# Folding scFv–Antigen Complexes at Scale

**DOI:** 10.64898/2026.07.01.730981

**Authors:** Ravi Shah, Jeffrey Ouyang-Zhang, Zachary Cohen, Maria Rosaria Briglia, Chi Zhang, Adam Klivans, Daniel Jesus Diaz

## Abstract

Accurate modeling of antibody–antigen (Ab–Ag) complexes is central to biologic development, yet the reliability and failures of modern Ab–Ag folding pipelines remain poorly characterized. Single-chain variable fragments (scFvs) are therapeutically important antibodies, but large-scale evaluations of structure prediction models on scFv–Ag complexes are largely lacking. We introduce a scalable bench-marking pipeline that generates large ensembles of scFv–Ag structure predictions by cofolding a curated subset of 3,800 Ab–Ag complexes from SAbDab using multiple state-of-the-art models under diverse inference-time settings. The resulting dataset, `SCALE` (**sc**Fv–**A**g Comp**L**ex **E**nsembles), includes standardized scFv–Ag sequences and around 200,000 predicted complexes spanning different models, sampling strategies, and auxiliary inputs. Using `SCALE`, we evaluate model performance in recovering correct scFv–Ag interfaces and assess the ability of existing confidence metrics to select the best structure from prediction ensembles. We find that while confidence scores effectively distinguish easy from hard scFv–Ag complexes, they often fail to identify the highest-quality interface for a given target. Further analysis shows that near-correct interfaces typically appear in ensembles but at low frequency, and inference-time choices such as sampling, recycling, and using evolutionary or structural information are crucial for accurate scFv–Ag complex predictions. Dataset and analysis code are available at https://huggingface.co/datasets/ravishah1/SCALE

## 1 Introduction

Antibodies underpin a large and rapidly growing class of therapeutics and diagnostics, enabling highly specific molecular recognition across oncology, immunology, and infectious disease (Scott et al., 2012; Murphy & Weaver, 2016; Burton & Hangartner, 2016). Central to their function is the formation of a correct antibody–antigen (Ab–Ag) complex: therapeutic efficacy and specificity depend on accurate epitope engagement and binding geometry. Reliable computational modeling of Ab–Ag complexes is therefore a critical component of modern antibody discovery pipelines, informing candidate screening, affinity optimization, and downstream design decisions.

Recent advances in protein structure prediction have enabled modeling of monomers and many protein complexes directly from sequence (Jumper et al., 2021; Abramson et al., 2024). However, Ab–Ag interactions remain a particularly challenging regime. Accurate Ab–Ag modeling requires resolving both highly flexible complementarity-determining regions (CDRs) and the global docking orientation between the paratope and the antigen’s epitope—errors in either can lead to poor interface quality even when individual chains are confidently folded. Single-chain variable fragments (scFvs) are a widely used antibody format in therapeutics, diagnostics, and display technologies, yet large-scale evaluations of structure prediction models on scFv–antigen (scFv–Ag) complexes are largely lacking.

In this work, we introduce `SCALE`, a scalable benchmarking framework for evaluating scFv–Ag cofolding at scale. Starting from 3,800 experimentally resolved Ab–Ag complexes, we construct standardized scFv–Ag sequence pairs and generate about 200,000 structure predictions with varying complex quality, using multiple state-of-the-art models and diverse inference-time settings. This ensemble-scale evaluation enables a systematic analysis of interface quality, sampling behavior, and the effectiveness of existing confidence metrics in identifying and filtering correct binding interfaces.

## 2 Related Work

### Folding Models

AlphaFold 2 is a protein structure predictor that takes an amino acid sequence as input and produces a full tertiary structure (Jumper et al., 2021). AlphaFold Multimer expands its capabilities to map a set of amino acid sequences into a full quaternary complex (Evans et al., 2021). AlphaFold 3 and its descendants (Abramson et al., 2024; Chai-Discovery et al., 2024; Passaro et al., 2025; Ouyang-Zhang et al., 2025) have extended this to other biomolecules while observing enhanced performance for antibody–antigen interfaces. Specialized antibody-focused models such as IgFold (Ruffolo et al., 2023) further improve monomer antibody structure prediction. (see Section A for inference-time folding settings)

### Interface Metrics

The DockQ metric (Basu & Wallner, 2016) is widely used to assess protein– protein interface (PPI) quality by combining the fraction of native contacts, interface RMSD, and ligand RMSD into a single composite score ranging from 0.0 to 1.0. DockQ is calibrated to align with the CAPRI benchmark (Janin et al., 2003), where scores of 0.0–0.23 indicate incorrect interfaces, 0.23–0.49 acceptable interfaces, 0.49–0.80 high-quality interfaces, and 0.80–1.0 near-native interfaces (see Section B for details on DockQ computation).

### Interface Confidence Models

Protein folding models are trained to produce auxiliary confidence scores. Metrics such as ipTM (Evans et al., 2021) aim to estimate the quality of protein–protein interfaces (PPIs) in predicted complexes. ipSAE (Dunbrack Jr, 2025) was introduced as a refinement of ipTM, with improved sensitivity to interface accuracy. pDockQ (Bryant et al., 2022) and pDockQ2 (Zhu et al., 2023) map folding-model confidence outputs to predicted DockQ. AbEpiScore (Clifford et al., 2025) is a learned neural network trained to predict Ab–Ag interface confidence.

### Benchmarking for Ab-Ag Structure Prediction

Recent studies have evaluated the performance of modern structure predictors on Ab–Ag complexes. Particularly Yin & Pierce (2024) benchmarks AlphaFold and AlphaFold Multimer on 427 non-redundant Ab–Ag complexes, analyzing interface accuracy. More recently, Hitawala & Gray (2025) investigate what AlphaFold 3 and related diffusion-based models learn about antibody and nanobody docking. However, these studies use a moderate scale and do not focus on scFv–Ag interfaces.

## 3 Method

In this section, we introduce our data pipeline which consists of: (i) a filtering stage of the SAbDab (Dunbar et al., 2014) database, where we aim to standardize a filtered subset of the data by creating scFv–Ag sequence pairs; (ii) to generate multiple structural predictions for each scFv–Ag complex, we fold the sequences under different folding models and inference settings; (iii) finally, we compute the DockQ score between the predicted and ground truth interfaces, aggregating additional measurements to benchmark the models.

### 3.1 Data Curation

#### SAbDab filtering

We *filter* the 10,133 PDB entries from SAbDab up to November 2025. We retain only samples with a single valid and annotated antigen chain, at least 16 residues long, and both valid and annotated antibody heavy and light chains. (Table 1)

**Table 1:** Dataset Curation Pipeline. Sequential filters applied to SAbDab to obtain the final dataset of 3,800 complexes.

| Filtering | Num. Complexes | Reduction |
| --- | --- | --- |
| SAbDab (raw) | 10,133 | — |
| Single Chain Ag Filter | 6,321 | -3,812 |
| $V_H/V_L$ Required Filter | 4,798 | -1,523 |
| Deduplicated Final Set | 3,800 | -998 |

#### scFv construction

For each filtered entry, we extract the amino acid sequences of the antigen chain, antibody heavy chain, and antibody light chain from the corresponding experimental structure. As shown in Figure 1a, we construct standardized scFv sequences by concatenating the antibody variable heavy domain (*V*_*H*_), a flexible glycine–serine linker, and the antibody variable light domain (*V*_*L*_): `scFv` = `VH` ∥ (`GGGGS`)_3_ ∥ `VL`, where *V*_*H*_ and *V*_*L*_ domains were identified using the ANARCI (Dunbar & Deane, 2016) tool under IMGT numbering. We then deduplicated samples with identical scFv–antigen sequence pairs. After filtering and scFv construction, the process produced 3,800 unique scFv–Ag sequence pairs.

**Figure 1:**
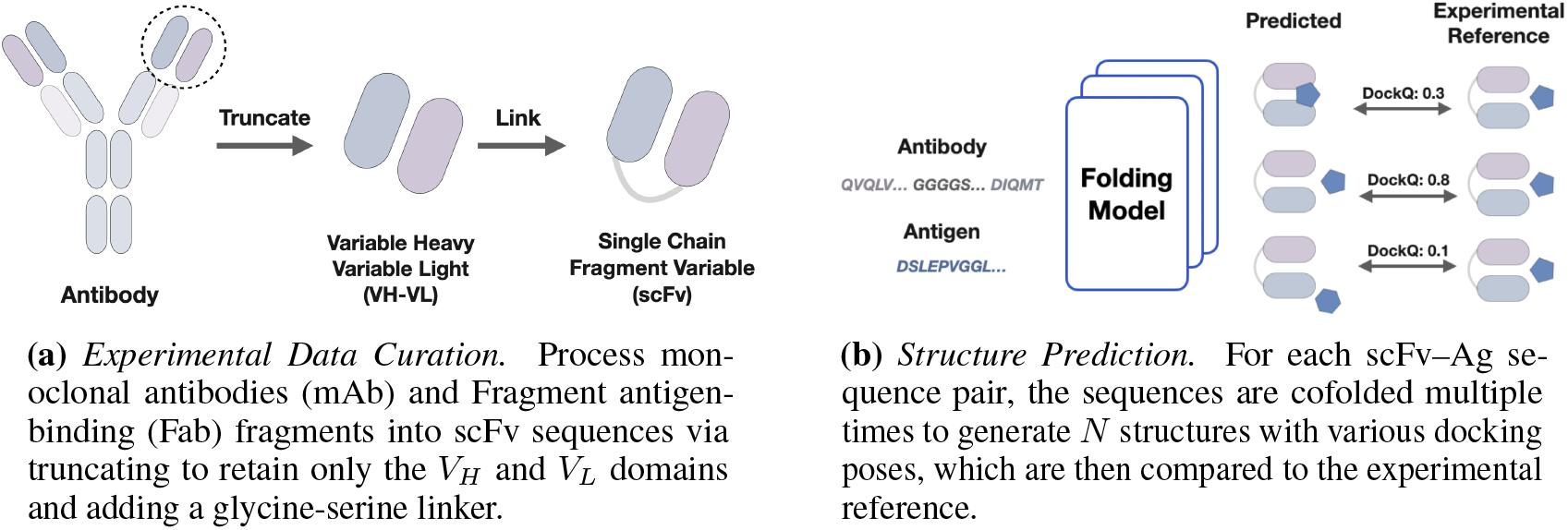
scFv–Antigen Dataset Generation Pipeline.

### 3.2 Structural Ensemble Pipeline

As shown in Figure 1b, starting from the filtered data, we cofold scFv-Ag sequence pairs to generate a diverse set of structural predictions. This involves simulating simple and challenging folding conditions for each sample. In this section, we outline details about the structure prediction pipelines. The folding configurations used to create our dataset are further detailed in Section D.

#### Backbone, Seed and Recycles Selection

We generate structures across multiple folding model backbones including AlphaFold 2.3 Multimer (Evans et al., 2021) via ColabFold (Mirdita et al., 2022), AlphaFold 3 (Abramson et al., 2024), Boltz-2 (Passaro et al., 2025), Chai-1 (Chai-Discovery et al., 2024), and Pairmixer (Ouyang-Zhang et al., 2025). Since the majority of the models are based on a diffusion process, we decide to introduce additional stochasticity in these modules by varying the inference seed. Using multiple randomized seeds help us increase the set of available structural predictions. Moreover, we adopt multiple settings for the *recycles* parameter. All models are run with 10 recycling steps, and to include lower-quality folds, we also generate AlphaFold 3 predictions with 1, 3, and 6 recycles. Further details on the composition of `SCALE` are provided in Table 3.

#### MSA and Template Generation

For each Ab–Ag complex, we construct antigen, scFv, and paired multiple sequence alignments (MSAs) using the ColabFold MSA pipeline (Mirdita et al., 2022) with the `uniref30_2302_db` database. The antigen MSA is always included to provide evolutionary context, while the scFv MSA is optionally included. We perform ablation experiments without the scFv MSA, motivated by the fact that antibody variable regions arise primarily from immune diversification rather than long-term evolutionary conservation. Structural templates are incorporated in a consistent manner for each complex by leveraging experimentally resolved structures from SAbDab. For each scFv–Ag complex, we retrieve the corresponding mAb or Fab structure from the RCSB and merge the two chains into a single continuous chain through systematic residue re-labeling and re-numbering. This representation aligns the template structure with the scFv-style binder format adopted in our pipeline, facilitating effective template matching during structure prediction.

### 3.3 Analysis Pipeline

Finally, we evaluate the quality of the generated samples using established structural metrics. For each filtered SAbDab complex, we generate the corresponding predicted scFv–Ag structures and compute the DockQ score (Basu & Wallner, 2016) between predicted and experimental structures using Open-Structure (Biasini et al., 2013). For benchmarking, we additionally run a variety of interface specific confidence models on the predicted structures including ipTM (Evans et al., 2021), ipSAE (Dunbrack Jr, 2025), pDockQ (Bryant et al., 2022), pDockQ2 (Zhu et al., 2023), and AbEpiScore (Clifford et al., 2025). Further implementation details in Section E.

## 4 Results

After constructing `SCALE`, we evaluated the ability of state-of-the-art cofolding models— AlphaFold 2.3, AlphaFold 3, Boltz-2, Chai-1, and Pairmixer—to generate high-quality scFv–Ag complex predictions. For each scFv–Ag complex in `SCALE`, we systematically varied random seeds, recycling depths, and the stochastic use of templates and MSAs (see Table 3) to generate complexes. This procedure produced 197,900 predicted complexes across five folding models.

We evaluated each predicted structure by computing its DockQ score with respect to the experimental SAbDab reference and report the resulting DockQ distributions for the *V*_*H*_ –Ag and *V*_*L*_–Ag interfaces in Figure 2a. Across the 3788, 59595, 22764, 3779, and 107974 predicted complexes, only 144, 8183, 6367, 475, 6854 achieved DockQ scores above 0.8, while 761, 22141, 12217, 1279, 27509 exceeded a DockQ threshold of 0.23 for AlphaFold 2.3, AlphaFold 3, Boltz-2, Chai-1, and Pairmixer, respectively. These results indicate that current state-of-the-art cofolding models fail to consistently produce high-quality scFv–Ag interfaces, even when supplied with multi-chain templates, MSAs, up to 10 recycling steps, and extensive seed sampling. Moreover, a substantial fraction of the evaluated complexes are present in the training sets of these models (Figure 5), underscoring their limited ability to generate realistic scFv–Ag complexes and accurately identify the antigen’s epitope that corresponds to a particular scFv paratope.

**Figure 2:**
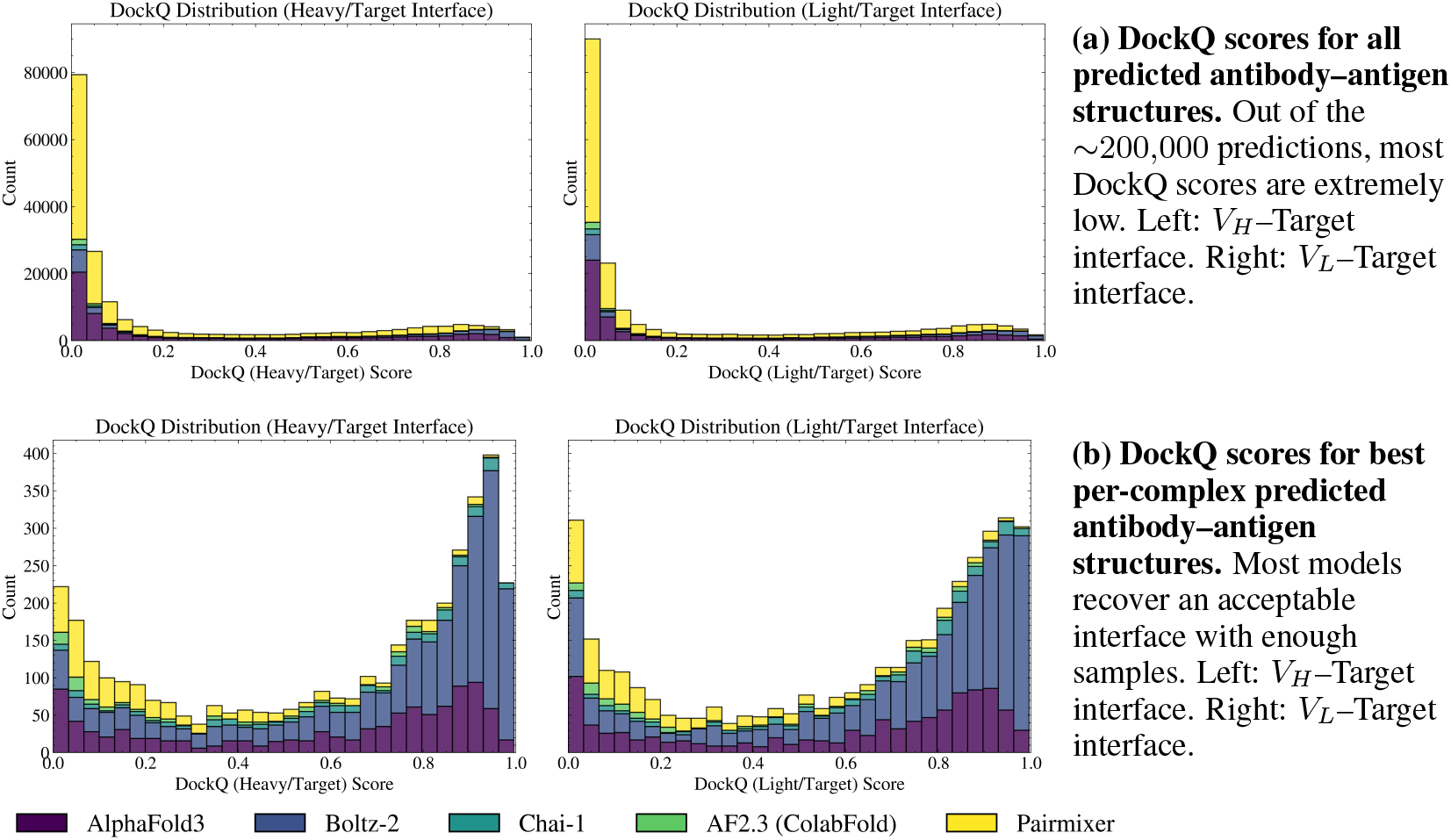
Docking Performance of scFv–antigen complexes. We evaluate multiple structure predictors on a curated split of SAbDab database with 3,800 antibody–antigen structures.

Next, we analyzed the highest-quality prediction per ensemble and report the corresponding *V*_*H*_ –Ag and *V*_*L*_–Ag DockQ scores in Figure 2b. The resulting distributions were bimodal, separating low- and high-quality cofolded complexes. Despite improved best-case performance, 879 of the 3800 scFv–antigen complexes fail to achieve a DockQ score above 0.23 under any evaluated setting, while only 256 consistently surpass this threshold across all conditions. For most targets, increasing the number of random seeds yields only marginal improvements (Figure 7). However, a small subset of complexes shows substantial gains from additional sampling, indicating that the effectiveness of inference-time settings is highly target-dependent. We further observe a strong coupling between *V*_*H*_ –Ag and *V*_*L*_–Ag interface quality (Pearson *r* = 0.958), with the *V*_*L*_–Ag interface being slightly more challenging to predict on average. Moving forward, unless otherwise specified, we therefore report DockQ as the mean of the *V*_*H*_ –Ag and *V*_*L*_–Ag interface scores.

Recent binder design pipelines (Pacesa et al., 2024; Mille-Fragoso et al., 2025; Stark et al., 2025) that have demonstrated experimental success rely heavily on AlphaFold confidence metrics and DockQ-inspired scores to filter and rank candidate structures. Therefore, we evaluated the ability of these confidence signals to correctly rank predicted scFv–Ag complexes (Table 2). Our analysis reveals a consistent and substantial gap between global ranking performance and per-ensemble discrimination across all evaluated metrics. While metrics such as ipTM, ipSAE, and pDockQ2 show strong global correlations with DockQ when predictions are pooled across all complexes, their average per-complex correlations and Top-*k* exact accuracies are significantly lower. This disparity indicates that existing confidence metrics are effective at coarse-grained discrimination—separating scFv–Ag complexes that are generally easy or difficult to cofold—but struggle to resolve fine-grained interface quality differences among alternative structures generated for the same complex. In practice, this limitation manifests as unreliable intra-complex reranking: confidence scores often fail to consistently identify the highest-quality PPI from large ensembles of candidate scFv–antigen structures. Taken together, these results suggest that while current confidence metrics are useful for global filtering between hard and easy scFv–antigen complexes, they are insufficient at consistently identifying the best-of-*N* PPI interface in high-throughput antibody–antigen cofolding pipelines.

**Table 2:** Evaluating metric performance on ranking predicted scFv-Ag complexes. We consider 3,298 SAbDab complexes with at least 50 successful folds, yielding a total of 174,563 predicted scFv-Ag structures. Global rank Spearman (*r*_*s*_) and Pearson (*r*_*p*_) correlations are computed by pooling all predicted scFv–Ag complexes across targets. Average correlations (avg *r*_*s*_, avg *r*_*p*_) are computed independently for each ensemble and then averaged across ensembles. Top-*k* measures the fraction of complexes for which the highest-quality prediction is ranked by the metric within the top *k* predictions.

| Method | $r_s$ (global) | $r_p$ (global) | avg $r_s$ | avg $r_p$ | Top-1 | Top-3 | Top-5 |
| --- | --- | --- | --- | --- | --- | --- | --- |
| ipTM (Evans et al., 2021) | 0.60 | 0.83 | 0.30 | 0.48 | 0.12 | 0.34 | 0.50 |
| ipSAE (Dunbrack Jr, 2025) | 0.71 | 0.82 | 0.32 | 0.49 | 0.11 | 0.32 | 0.47 |
| pDockQ (Bryant et al., 2022) | 0.32 | 0.49 | 0.21 | 0.35 | 0.10 | 0.27 | 0.41 |
| pDockQ2 (Zhu et al., 2023) | 0.74 | 0.79 | 0.31 | 0.47 | 0.13 | 0.34 | 0.50 |
| AbEpiScore (Clifford et al., 2025) | 0.62 | 0.70 | 0.22 | 0.33 | 0.07 | 0.19 | 0.28 |

**Table 3:**
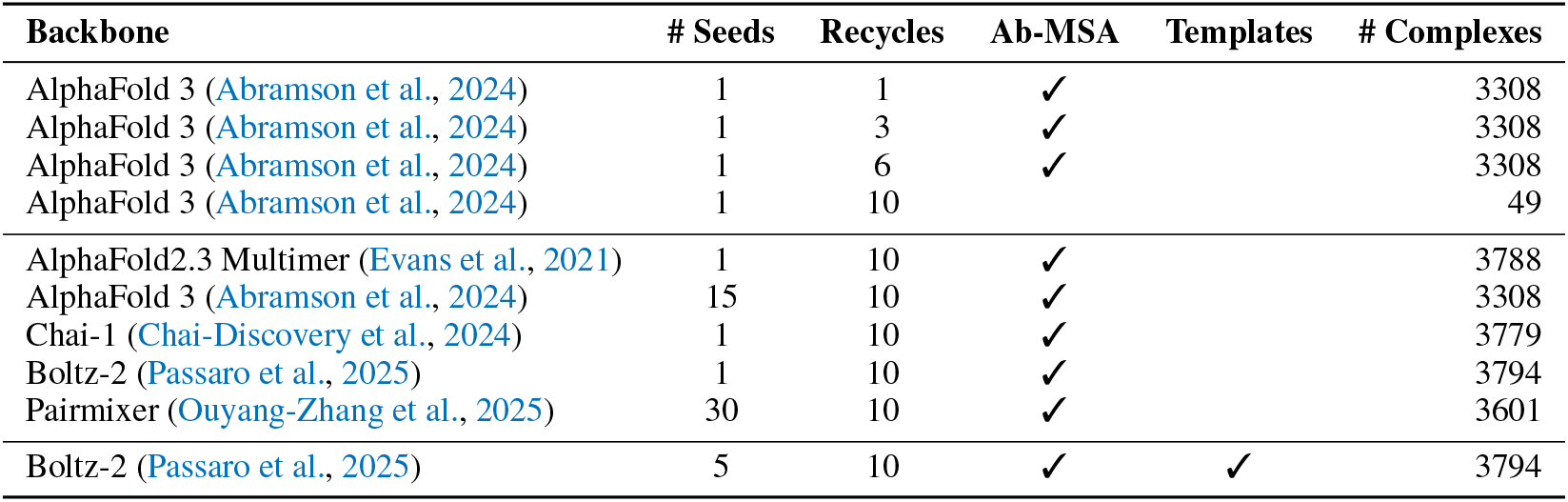
Folding configurations. Summary of model backbones and inference-time settings used to generate structure predictions for benchmarking.

Several consistent patterns emerge across inference settings. First, high single-chain confidence does not imply accurate quaternary complex formation (shown in Figure 11). Most predictions achieve high pLDDT scores yet exhibit poor PPI quality, revealing a decoupling between tertiary and quaternary structural accuracy. Second, increasing inference-time refinement primarily improves best-case rather than average complex accuracy. Additional recycling steps occasionally yield near-native interfaces, but the median DockQ score remains largely unchanged (Figure 8). Third, auxiliary input influences model behavior—supplying an antibody MSA increases the likelihood of sampling high-quality interfaces, despite the limited biological relevance of MSAs for antibodies (Figure 9). Finally, certain physical properties of scFv–Ag interfaces show little correlation with interface quality (Figure 12). However, other PyRosetta (Chaudhury et al., 2010) derived physics based properties— such as interface shape complementarity and estimated binding energy—are roughly as effective as commonly used confidence metrics (e.g., ipTM) for ranking predicted structures, in terms of both global and per-complex Spearman correlation (Table 4).

**Table 4:** Rosetta-derived feature correlations grouped by physical interaction type. Spearman correlation (*r*_*s*_) is reported globally across all structures and averaged per ensemble (per-PDB).

| Feature | Global $r_s$ | Per-PDB $r_s$ |
| --- | --- | --- |
| *Interface Geometry & Binding Strength (shape, affinity, size)* |  |  |
| interface_shape_complementary | 0.60 | 0.26 |
| interface_dG | -0.57 | -0.27 |
| interface_dG_SASA_ratio | -0.57 | -0.28 |
| interface_dSASA | -0.26 | -0.06 |
| interface_fraction | -0.27 | -0.10 |
| interface_nres | -0.13 | -0.03 |
| interface_packstat | 0.19 | -0.02 |
| *Steric Repulsion* |  |  |
| interface_fa_rep | -0.55 | -0.25 |
| complex_fa_rep | -0.55 | -0.25 |
| target_fa_rep | -0.54 | -0.24 |
| binder_fa_rep | -0.49 | -0.24 |
| *Van der Waals Attraction* |  |  |
| interface_fa_atr | 0.27 | 0.08 |
| complex_fa_atr | 0.42 | 0.20 |
| target_fa_atr | 0.41 | 0.18 |
| binder_fa_atr | 0.27 | 0.21 |
| *Solvation (hydrophobic burial)* |  |  |
| interface_fa_sol | -0.28 | -0.09 |
| complex_fa_sol | -0.42 | -0.19 |
| target_fa_sol | -0.41 | -0.16 |
| binder_fa_sol | -0.20 | -0.20 |
| interface_hydrophobicity | 0.04 | 0.03 |
| surface_hydrophobicity | -0.01 | 0.08 |
| *Electrostatics* |  |  |
| interface_fa_elec | -0.27 | -0.12 |
| complex_fa_elec | 0.37 | -0.07 |
| target_fa_elec | 0.38 | -0.07 |
| binder_fa_elec | -0.15 | -0.08 |
| *Hydrogen Bonding* |  |  |
| interface_hbond_percentage | 0.46 | 0.22 |
| interface_interface_hbonds | 0.41 | 0.21 |
| complex_hbond_bb_sc | 0.30 | -0.02 |
| complex_hbond_sc | 0.22 | -0.13 |
| target_hbond_bb_sc | 0.32 | -0.03 |
| target_hbond_sc | 0.27 | -0.13 |
| binder_hbond_bb_sc | -0.10 | 0.00 |
| binder_hbond_sc | -0.09 | -0.08 |
| *Unsatisfied Polar Penalties (buried unsatisfied H-bonds)* |  |  |
| interface_delta_unsat_hbonds | -0.36 | -0.14 |
| interface_delta_unsat_hbonds_percentage | -0.35 | -0.15 |
| *Sidechain Conformation (rotamer strain)* |  |  |
| complex_fa_dun | -0.36 | 0.20 |
| target_fa_dun | -0.39 | 0.20 |
| binder_fa_dun | 0.14 | 0.18 |
| *Anisotropic Solvation / Water-mediated* |  |  |
| interface_lk_ball_wtd | -0.25 | -0.09 |
| complex_lk_ball_wtd | 0.35 | 0.02 |
| target_lk_ball_wtd | 0.37 | 0.02 |
| binder_lk_ball_wtd | -0.10 | 0.02 |
| *Global Energy* |  |  |
| binder_score | -0.51 | -0.24 |

## 5 Discussion

This work provides a large-scale evaluation of modern structure cofolding models on scFv–Ag complexes by folding under varying inference settings to create an ensemble of structures per complex. Near-correct interfaces typically appear in ensembles but at low frequency, implying a sampling and selection bottleneck rather than representational capacity alone. A key challenge exposed by our results is the inability of existing confidence metrics to reliably select high-quality interfaces on a per-complex basis. Although scores such as ipTM, ipSAE, and pDockQ2 correlate well with interface quality when predictions are pooled globally, they perform poorly at identifying the best structure within large ensembles. Notably, certain physics-based features describing the interface are just as effective as confidence metrics for selecting high-quality interfaces. In terms of producing high-quality quaternary structures, we found that inference-time choices often impact best-case performance but do not consistently improve typical outcomes.

Overall, our results suggest that (i) accurate scFv–Ag interface prediction remains challenging for modern folding models; (ii) with sufficient sampling and diverse inference settings, high-quality interfaces often emerge; (iii) reliable structure re-ranking is therefore a critical bottleneck, yet existing confidence metrics perform poorly at this task; and (iv) physics-based descriptors of the interface should be considered with confidence metrics for evaluating docking quality of a complex.

## 6 ACKNOWLEDGMENT

This work is supported by the NSF AI Institute for Foundations of Machine Learning (IFML) and UT-Austin Center for Generative AI.

## A Structure Prediction Sampling Settings

We detail the sampling parameters below.

**Random seeds** control the stochasticity of diffusion-based structure prediction by initializing the noise sampling process. Changing the seed alters the diffusion trajectory while keeping all inputs fixed, sometimes producing substantially different final structures. Sampling multiple seeds is a useful tool for exploring structural uncertainty and generating diverse predictions.

**Recycling** refers to iteratively feeding intermediate structure predictions back into the model to refine internal representations and improve geometric consistency. Recycling is specified as an integer hyperparameter and typically improves interface quality at the cost of increased inference time and compute.

**Multiple sequence alignments (MSAs)** provide evolutionary context by encoding residue conservation and co-evolutionary signals across homologous sequences. MSAs are supplied as alignment files (e.g., `.a3m`) and often substantially improve the tertiary structure of the global fold.

**Structural templates** supply an explicit structural prior in the form of experimentally determined coordinates from related proteins. Templates are passed to the model as structure files (e.g., PDB or CIF) and can guide folding toward biologically plausible conformations.

## B Interface Quality Metrics

The DockQ metric (Basu & Wallner, 2016) is a standard interface quality measure used in the protein–protein docking community to quantify agreement between predicted and experimentally determined complexes. DockQ integrates three established interface metrics—FNAT, iRMSD, and LRMSD—into a single continuous score.

**Fraction of Native Contacts (FNAT)** measures the recovery of native inter-chain contacts in a predicted complex. Let *C*_native_ denote the set of inter-chain residue contacts in the experimentally determined reference structure and *C*_pred_ the corresponding set in the predicted structure. A contact is defined by a heavy-atom distance threshold between residues across chains (typically 4–5 Å) (Janin et al., 2003; Basu & Wallner, 2016). FNAT is defined as

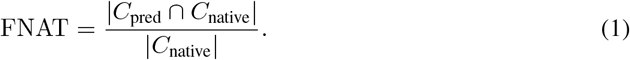

**Interface RMSD (iRMSD)** measures the root mean squared deviation over interface residues after optimal rigid-body superposition of the two complexes. Interface residues are defined as those participating in inter-chain contacts in the native structure, and superposition is performed over the full complex prior to computing RMSD on the interface subset (Janin et al., 2003; Basu & Wallner, 2016).

**Ligand RMSD (LRMSD)** measures the RMSD of the smaller (ligand) chain after aligning the larger (receptor) chain to the reference, capturing global rigid-body docking accuracy and relative chain placement (Janin et al., 2003; Basu & Wallner, 2016).

**DockQ labeling function** integrates FNAT, iRMSD, and LRMSD, unifying interface contact recovery and geometric accuracy into a single continuous, interpretable score. Formally,

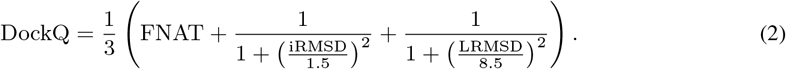

DockQ is designed to correlate with CAPRI docking quality categories (Basu & Wallner, 2016). DockQ scores are evaluated using four ranges: 0.00–0.23, 0.23–0.49, 0.49–0.80, and 0.80–1.00. These correspond to poor, acceptable, high-quality, and near-native interfaces, respectively.

## C SAbDab Filtering and scFv Dataset Construction

We construct our scFv–antigen dataset through a multi-stage filtering and processing pipeline based on SAbDab metadata and experimentally resolved structures. The full procedure is outlined below.

### Load and standardize SAbDab metadata

We begin from the full SAbDab summary table, which contains per-entry metadata including PDB identifiers, chain annotations, experimental method, resolution, and antibody format. The initial dataset contains 10133 total SAbDab entries (PDB files).

### Antigen and chain-level filtering

We first remove entries without an annotated antigen chain. We then restrict to complexes containing a single antigen chain, and explicitly exclude unreliable samples in which the antigen chain sequence is identical to a heavy or light chain sequence. After this step, 6321 entries remain.

### Antibody format and chain requirements

We exclude pre-annotated scFv entries and retain only complexes containing both a heavy and a light chain, corresponding to full-length antibodies (mAbs, monoclonal antibodies) or Fab fragments (fragment antigen-binding). Entries are required to contain both a valid heavy chain annotation and a valid light chain annotation. After this step 4798 entries remain.

### Structure loading and sequence extraction

For each metadata entry, we load the corresponding experimental structure and extract amino acid sequence for the antigen chain and antibody heavy and light chains using Biopython (Cock et al., 2009).

### scFv construction

For each heavy–light antibody, we construct a single-chain variable fragment (scFv) by concatenating the variable heavy (*V*_*H*_) and variable light (*V*_*L*_) domains using a flexible glycine–serine linker (`(GGGGS)`_3_). Domain boundaries and framework/CDR regions are identified during construction and stored explicitly using ANARCI (Dunbar & Deane, 2016).

### Dataset assembly and deduplication

Each processed complex is represented as a row containing the scFv sequence, antigen sequence, chain identifiers, and per-region length metadata. We deduplicate identical scFv–antigen sequence pairs, and discard targets shorter than 16 residues. The final dataset contains 3800 entries.

## D Dataset Generation & Folding Configurations

Table 3 summarizes the folding model backbones and inference-time configurations used to generate the benchmarking dataset. In total, we generated 197900 predicted structures across 3800 unique scFv– antigen sequence pairs. These predictions span five distinct folding backbones and ten configurations that vary in the number of stochastic seeds, recycling steps, inclusion of antibody MSAs (Ab–MSA), and use of structural templates.

To probe the sensitivity of scFv–antigen folding to inference-time choices, we include configurations designed to increase task difficulty by limiting model refinement or auxiliary information. Specifically, we evaluate AlphaFold 3 under reduced recycling budgets and with the antibody MSA removed (Table 3, top section). These settings test the robustness of interface prediction when refinement or evolutionary context is restricted.

For standardized comparison across folding backbones, we additionally evaluate each model using a common inference configuration consisting of 10 recycling steps and inclusion of the Ab–MSA (Table 3, middle section). This allows direct comparison of interface quality across AlphaFold 2.3 Multimer, AlphaFold 3, Boltz-2, Chai-1, and Pairmixer under matched conditions. Pairmixer is run with an increased number of stochastic seeds to exploit its computational efficiency and assess the impact of extensive sampling.

Finally, we include a small number of configurations that provide structural templates derived from experimentally resolved complexes (Table 3, bottom section). While these templates reveal some ground-truth structural information, they serve as an upper bound on achievable interface quality and allow us to assess how strongly template guidance influences scFv–antigen docking performance.

Not all configurations are applied to every scFv–antigen pair. In particular, some settings are restricted to the 49 complexes comprising the AlphaFold 3 antibody–antigen test set (Abramson et al., 2024). For the remaining configurations, we attempt to fold all 3800 complexes; however, only a subset are successfully completed due to MSA compatibility constraints and practical computational limitations.

## E Implementation Details of Existing Confidence Scores

We implemented the existing confidence scores using the definitions and specific details that follow.

### E.1 Interface predicted TM-SCORE (IPTM)

Let *C*_1_ and *C*_2_ denote two distinct chains, and let PAE_*ij*_ be the predicted aligned error (in Å) between residues *i* and *j*. Our PAE matrix was sliced to exclude all linker indices.

For an isolated two-chain system, define

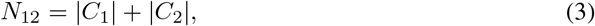

and compute a single normalization constant

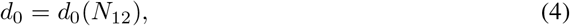

where *d*_0_(·) follows the standard TM-score length normalization.

Define the inter-chain PAE blocks

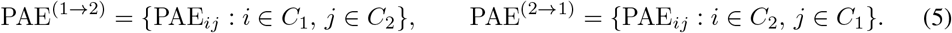

For residue *i* ∈ *C*_1_, define

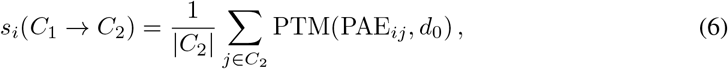

and analogously for *C*_2_ → *C*_1_. The directional scores are

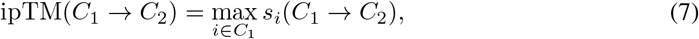

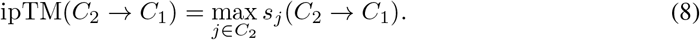

The final interface predicted TM-score is

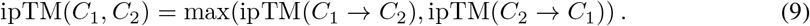

It should be noted that this ipTM score will differ slightly from AlphaFold’s ipTM. AlphaFold computes a probability-weighted sum on PTM transformations of PAE possibilities, while our PAE values are already compressed into a single value via probability-weighted sums and passed into the PTM function. Nonetheless, our ipTM has a 0.959 spearman correlation to the ipTM directly outputted from AF3 as shown in Figure 3. Chain-specific ipTM scores (*V*_*H*_ –target and *V*_*L*_–target) are computed by masking the PAE matrix to include only residue pairs belonging to the corresponding chains.

**Figure 3:**
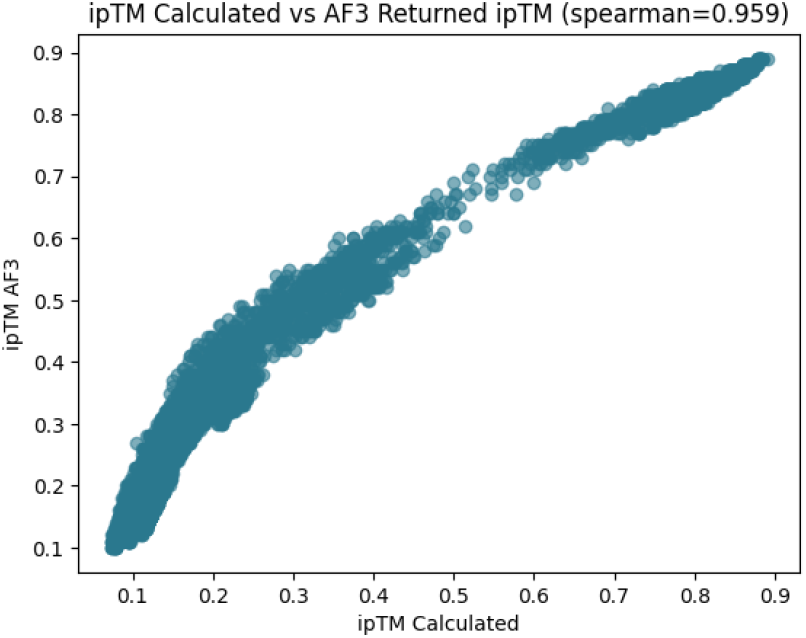
iPTM validation. Comparison between our computed ipTM score and the ipTM value reported directly by AlphaFold 3. We must perform our own ipTM calculation to generate scores for the *V*_*H*_ –target and *V*_*L*_–target interfaces from the scFv–Ag complex.

### E.2 Interface predicted Structural Alignment Error (IPSAE)

ipSAE is defined as a score between two chains derived directly from the PAE matrix. Our PAE matrix was sliced to exclude linker residues. Let *C*_1_ and *C*_2_ denote two distinct chains, and let PAE_*ij*_ be the predicted aligned error (in Å) between residue *i* ∈ *C*_1_ and residue *j* ∈ *C*_2_. For each residue *i* ∈ *C*_1_, define

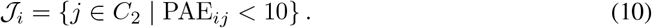

Let

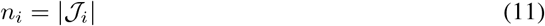

be the number of interface residues associated with residue *i*. Residues with *n*_*i*_ = 0 are excluded. For each valid residue *i*, define a per-residue normalization constant

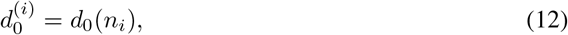

where *d*_0_ follows the standard TM-score length normalization. The residue-level interface score is

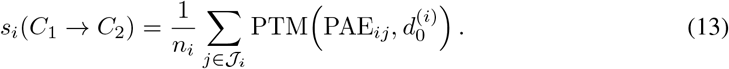

The predicted structural alignment error of the directional interface is defined as

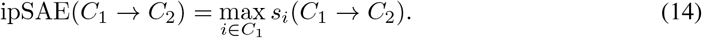

For any chain pair (*C*_1_, *C*_2_), we define a final symmetric interface score as

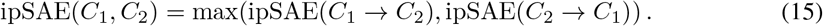

Chain-specific ipSAE scores (*V*_*H*_ –target and *V*_*L*_–target) are computed by masking the PAE matrix to include only residue pairs belonging to the corresponding chains.

### E.3 Predicted DockQ (PDockQ)

Let *C*_1_ and *C*_2_ denote two distinct chains. Residue *i* has Cartesian coordinates **x**_*i*_ ∈ ℝ^3^ and confidence score pLDDT_*i*_. The pLDDT matrix was sliced to exclude linker residues.

#### Interface contacts

For residues *i* ∈ *C*_1_ and *j* ∈ *C*_2_, define

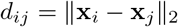

Residues are considered in contact if

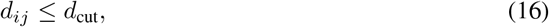

with *d*_cut_ = 8 Å. Let *C*_12_ be the set of all inter-chain contacts and

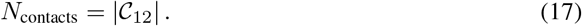

If *N*_contacts_ = 0, the pDockQ score is set to zero. Let

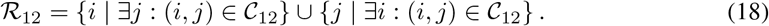

The mean interface confidence is

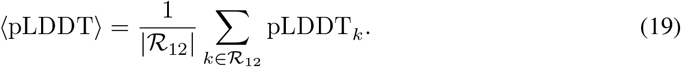

Define

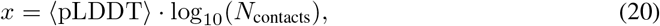

and compute

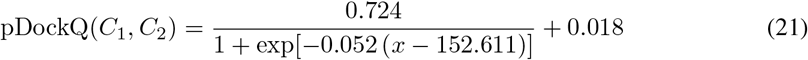

Chain-specific pDockQ scores (*V*_*H*_ –target and *V*_*L*_–target) are computed by masking the pLDDT vector and inter-chain contact map to include only residue pairs belonging to the corresponding chains.

### E.4 Predicted DockQ2 (PDockQ2)

Let *C*_1_ and *C*_2_ denote two distinct chains. Residue *i* has coordinates **x**_*i*_ ∈ ℝ^3^ and confidence pLDDT_*i*_. Let PAE_*ij*_ be the predicted aligned error (in Å) between residues *i* and *j*. Our PAE and pLDDT matrices are sliced to exclude linker residues.

Define

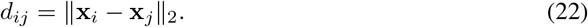

A pair (*i, j*) with *i* ∈ *C*_1_, *j* ∈ *C*_2_ is a contact if

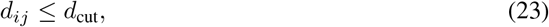

with *d*_cut_ = 8 Å. Let *C*_12_ denote the set of all such inter-chain contacts. If |*C*_12_| = 0, we set pDockQ2(*C*_1_, *C*_2_) = 0.

For each contacting pair (*i, j*) ∈ *C*_12_, define

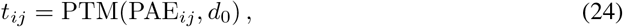

where *d*_0_ is fixed (default *d*_0_ = 10). The mean contact score is

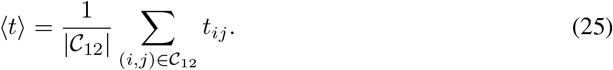

Let ℛ_12_ be the set of residues participating in at least one contact:

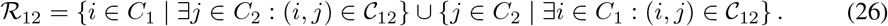

Then

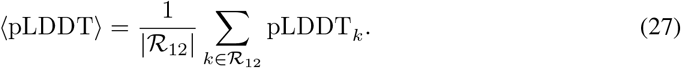

Define

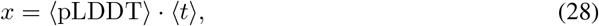

and compute

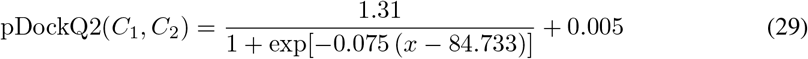

Chain-specific pDockQ scores (*V*_*H*_ –target and *V*_*L*_–target) are computed by masking the PAE matrix, pLDDT vector, and inter-chain contact map to include only residue pairs belonging to the corresponding chains.

### E.5 AbEpiScore

We implement AbEpiScore using the default, official implementation codebase: https://github.com/mnielLab/AbEpiTope-1.0/tree/main.

## F Additional Results

### Dataset distribution

As shown in Figure 4, the majority of predictions across all models yield low DockQ scores, indicating generally poor interface quality. However, a small fraction of predictions still achieve near-native interfaces, suggesting that correct binding modes are occasionally recovered.

**Figure 4:**
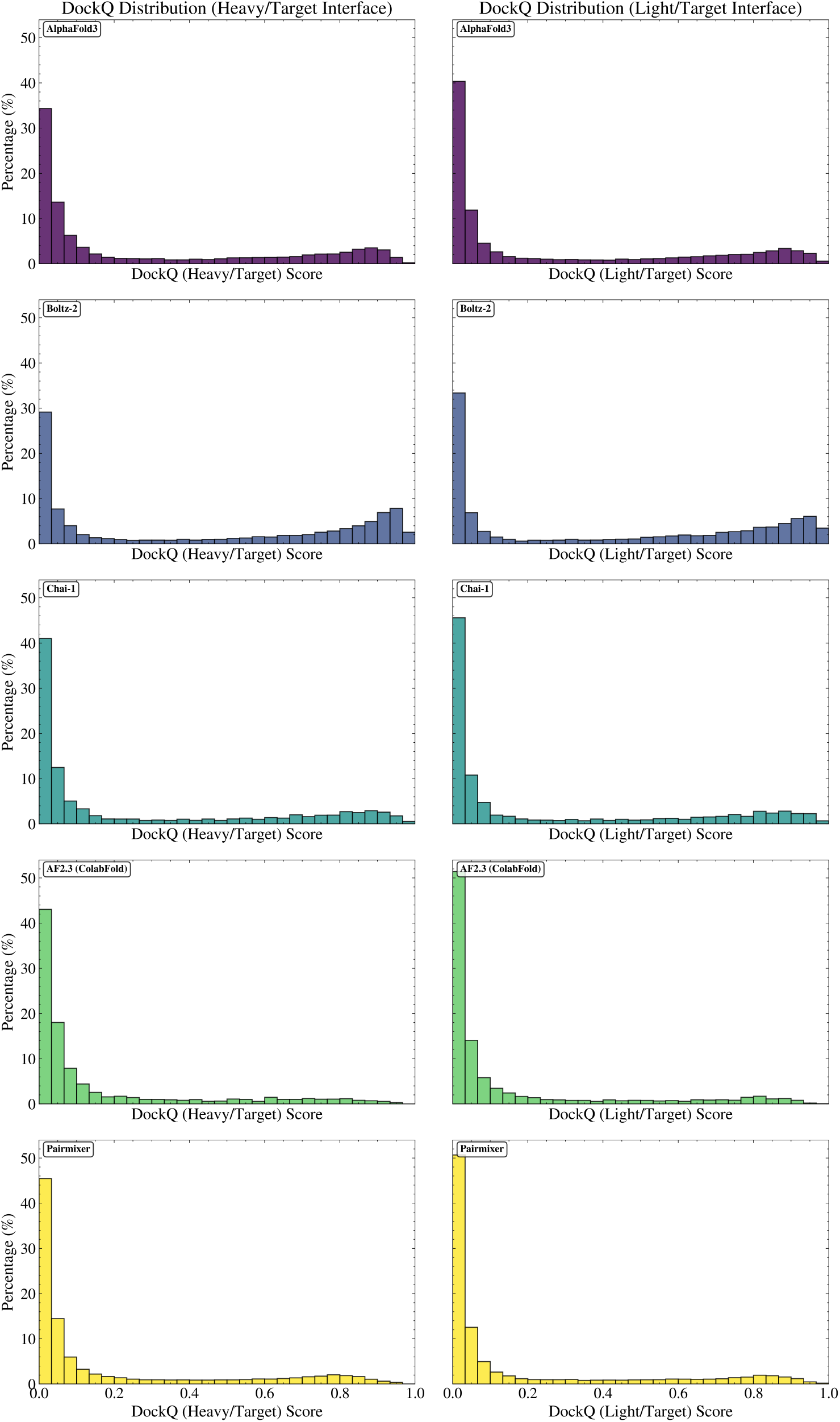
DockQ distribution across folding models. Distributions of DockQ scores for *V*_*H*_ –target (left) and *V*_*L*_–target (right) interfaces, stratified by folding backbone. Predicted structures from all folding configurations specified in Table 3 are included.

Boltz-2 appears to exhibit improved performance relative to other models. However, Figure 5 suggests that this improvement is likely attributable to the later training cutoff date, which increases the likelihood that some complexes were seen during training. Consistent with this, complexes released after the training cutoff (unseen by the model) typically show degraded performance. Notably, even among complexes within the training set, many are not accurately predicted, indicating that memorization alone does not fully explain model performance.

**Figure 5:**
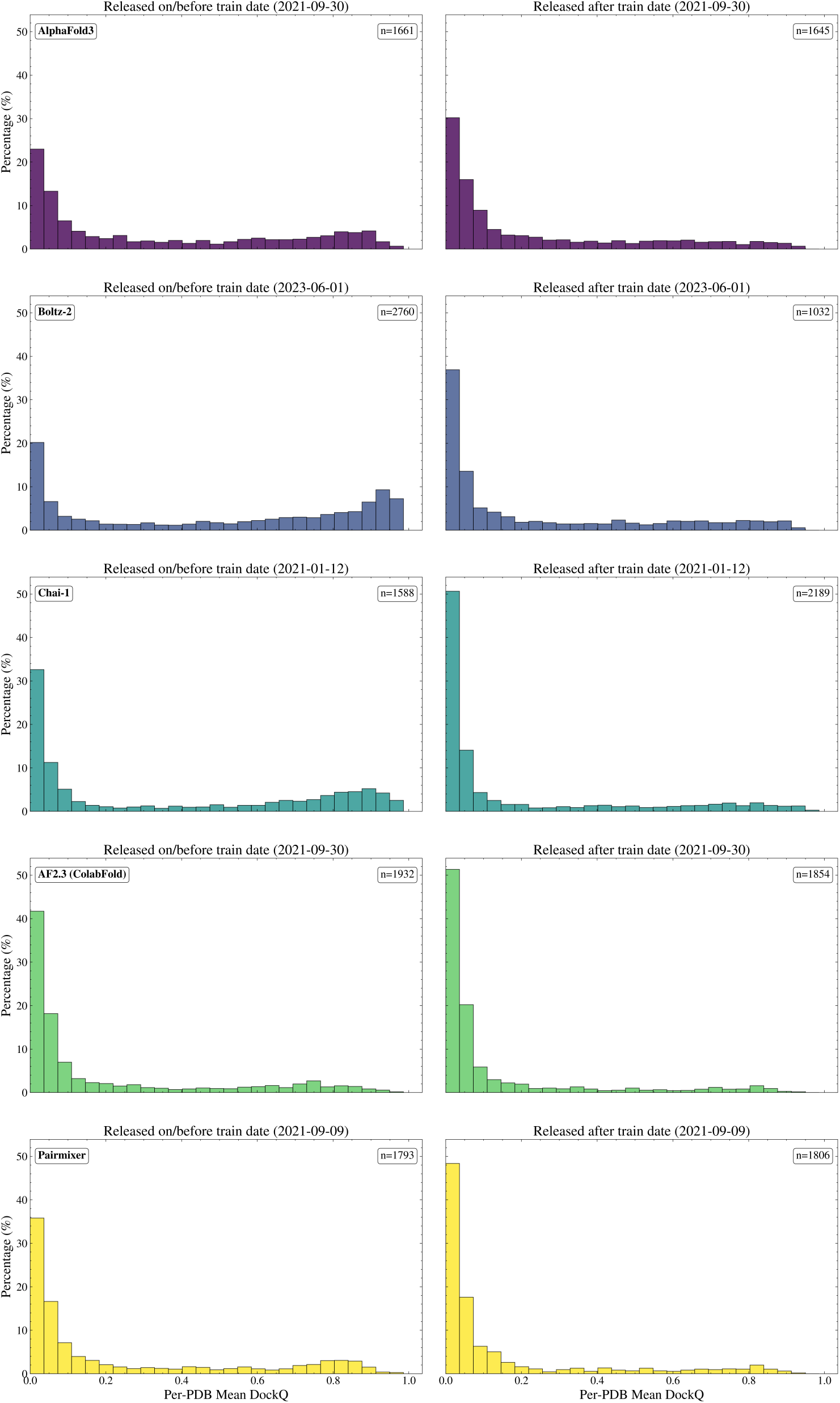
DockQ distribution before and after training date cutoff. Distributions of mean per-ensemble DockQ scores, stratified by folding backbone. Left: PDBs released on or before the model-specific training cutoff (seen during training). Right: PDBs released after the cutoff (unseen during training).

### Impact of inference-time settings

We find that the impact of inference-time parameters is highly dependent on the specific complex being predicted. As shown in Figure 6 and Figure 7, most complexes exhibit low variability in DockQ across stochastic samples, indicating that repeated inference typically converges to similar docking poses. However, for a smaller subset of targets, sampling additional seeds can uncover substantially different and higher-quality interfaces. Similarly, Figure 8 shows that increasing the number of recycling steps primarily improves already accurate predictions, occasionally refining them from acceptable or high-quality to near-native interfaces, while having limited effect on typical performance.

**Figure 6:**
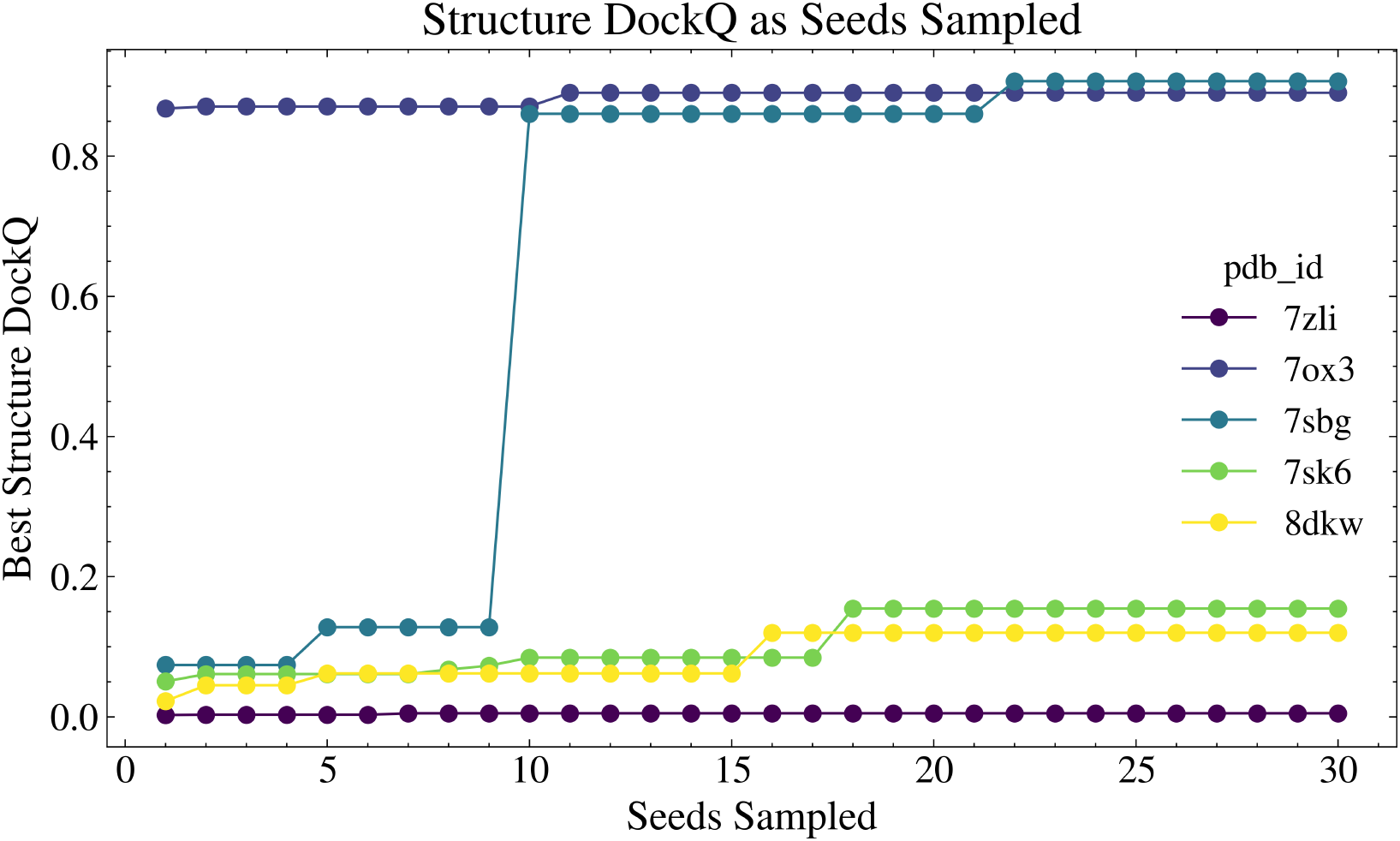
Effect of stochastic sampling on interface recovery. Best DockQ score as a function of the number of Pairmixer (Ouyang-Zhang et al., 2025) random seeds sampled, shown for five representative complexes from the AlphaFold 3 antibody–antigen test set (Abramson et al., 2024). Each curve reports the maximum DockQ observed among the first *N* sampled seeds. For most complexes, increasing the number of seeds yields only marginal improvements, indicating diminishing returns from additional stochastic sampling. However, for a subset of targets, additional sampling can uncover substantially higher-quality interfaces that are not observed at low sample counts.

**Figure 7:**
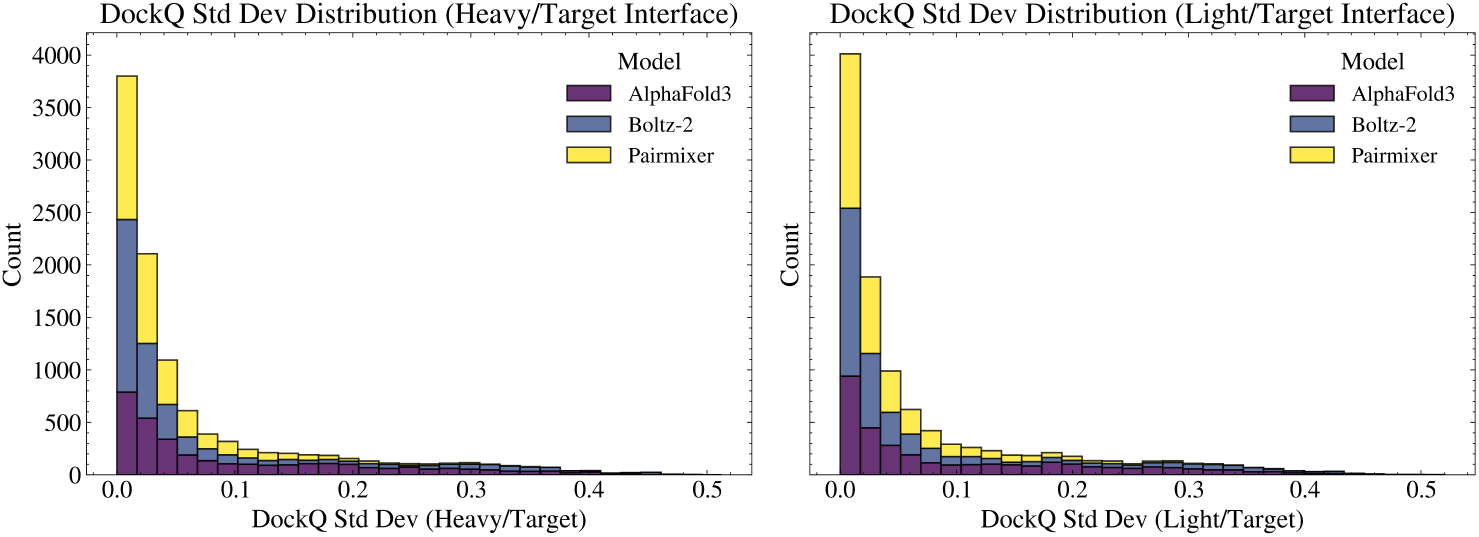
Seed-induced interface diversity across folding models. Distribution of per-ensemble standard deviation of DockQ across random seeds for *V*_*H*_ –target (left) and *V*_*L*_–target (right) interfaces, stratified by model. Most complexes exhibit low variance, indicating that repeated stochastic inference produces similar interface qualities. A smaller subset shows substantial variability, reflecting cases where different seeds explore distinct docking modes. This suggests that while seed ensembling can increase diversity, many antibody–antigen complexes remain stably confined to similar docking poses.

**Figure 8:**
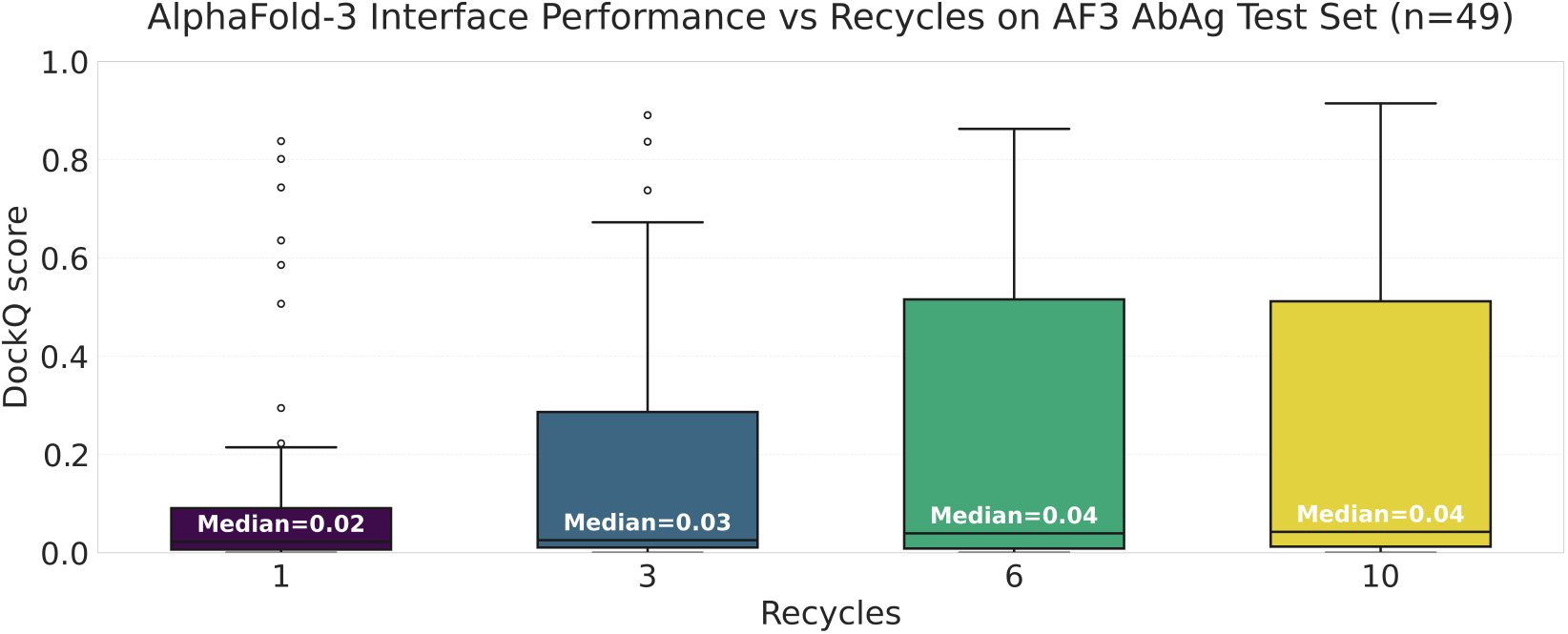
Recycling improves best-case but not typical interface quality. Distribution of DockQ scores produced by AlphaFold 3 on the antibody–antigen test set (*n* = 49 complexes) under varying numbers of recycling steps. While increasing the number of recycles enables substantially better top-end predictions, including occasional near-native interfaces, the median DockQ remains low and changes only marginally across settings. This suggests that recycling primarily increases variability across predictions, improving best-case outcomes without consistently improving typical performance.

Figure 9 demonstrates that providing an antibody MSA is consistently beneficial: on the AlphaFold 3 test set, incorporating MSAs reduces the fraction of low-quality predictions (DockQ *<* 0.23) by approximately 22%. In contrast, providing structural templates yields only modest improvements, suggesting limited impact on quaternary structure accuracy (Figure 10). Regardless of the inference settings used, high-quality tertiary structure is not predictive of quaternary structure accuracy, as illustrated in Figure 11.

**Figure 9:**
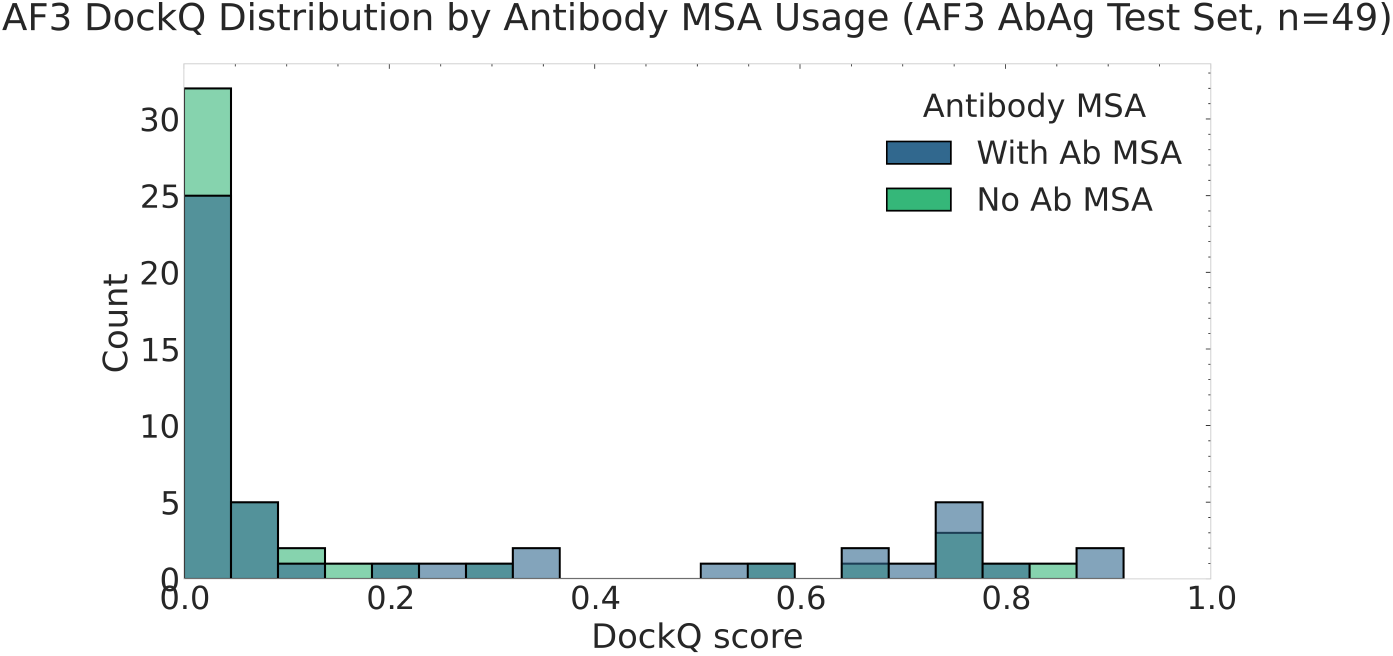
Antibody MSAs improve interface quality. Distribution of DockQ scores produced by AlphaFold 3 on the antibody–antigen test set (*n* = 49 complexes), comparing predictions generated with (blue) and without (green) an antibody MSA. Including an antibody MSA increases the frequency of high-quality interface predictions, while removing it leads to a shift toward low DockQ scores. This trend suggests that antibody MSAs provide a strong inductive bias that stabilizes predictions, even if their direct biological relevance for antibody sequences is limited.

**Figure 10:**
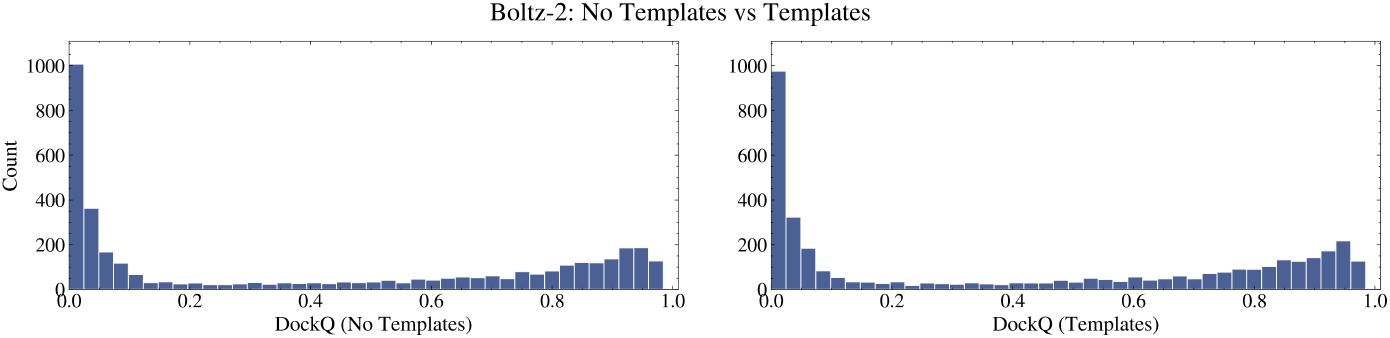
Limited impact of structural templates in Boltz-2. Distribution of DockQ scores produced by Boltz-2 without (left) and with (right) structural templates. The distributions are broadly similar, indicating that template information provides only modest improvements in interface quality for scFv–Ag docking.

**Figure 11:**
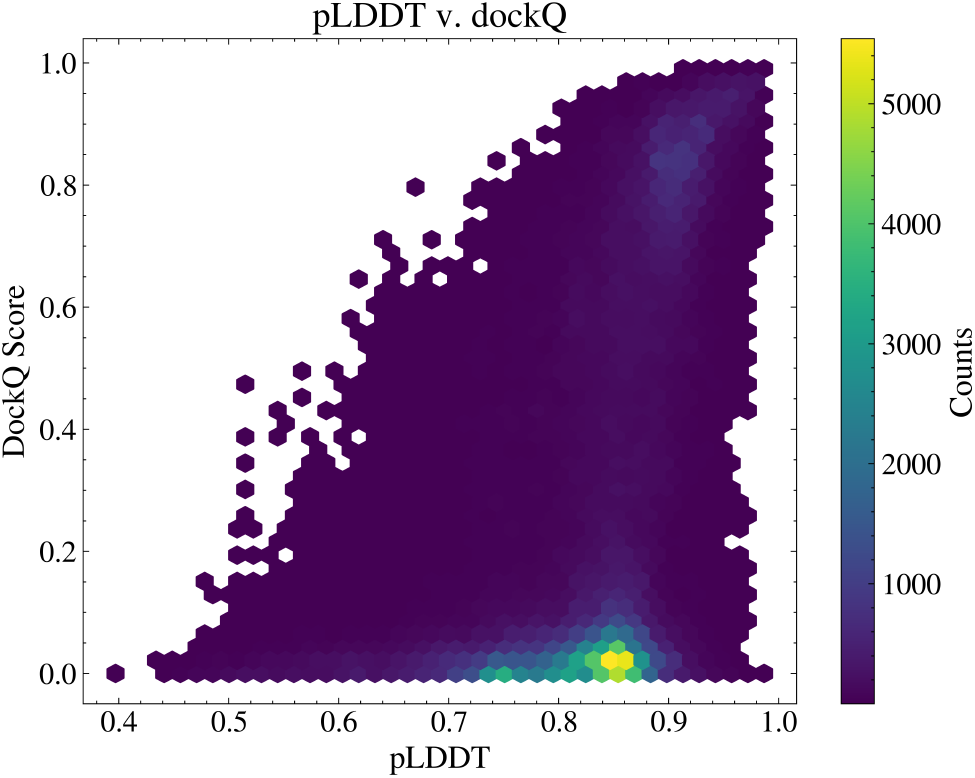
High pLDDT does not imply correct scFv–Ag interfaces. Hexbin density plot of predicted local distance difference test (pLDDT) versus DockQ across 197900 predicted scFv-Ag complexes across the entire dataset. Over 75% of predictions have mean pLDDT above 0.8, indicating that modern folding models consistently recover confident tertiary structures. However, a substantial fraction of these high-pLDDT predictions still achieve very low DockQ scores, reflecting incorrect binding orientations or epitope placement. This decoupling highlights a key challenge in scFv–Ag modeling: accurate single-chain folding does not guarantee correct quaternary structure or high-quality scFv–Ag interfaces.

### Secondary structure analysis

For each complex, we analyze secondary structure of the epitope. We define the epitope as any antigen residues within 8 Å of the antibody heavy or light chain in the experimental structure. Secondary-structure assignments are obtained using STRIDE (Frishman & Argos, 1995), from which we compute the fraction of helix, *β*-strand, and coil residues for each epitope. We then examine the correlation between the secondary structure of a complex’s epitope and the mean predicted DockQ of that structure across its ensemble of folds. As shown in Figure 12, secondary structure is a weak indicator of whether an interface will be easy or difficult for the model to predict.

**Figure 12:**
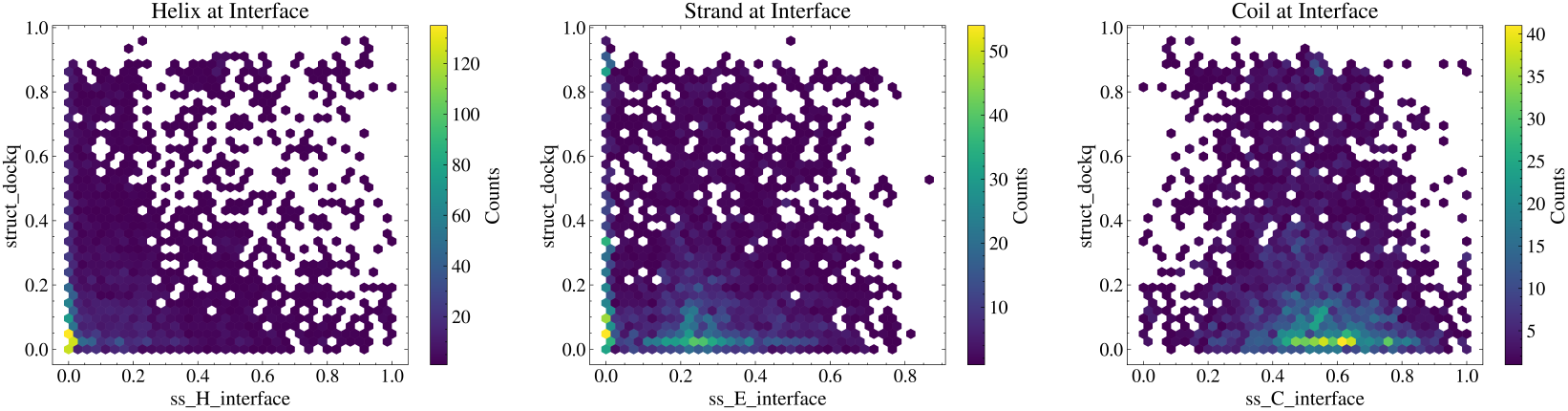
Epitope secondary structure is weakly predictive of interface quality. Hexbin density plots of DockQ versus the fraction of helix (left), *β*-sheet (center), and coil (right) residues within the antigen epitope, defined as residues within 8 Å of the antibody in the experimental complex. All three secondary-structure classes show weak or negligible correlation with DockQ, indicating that both high- and low-quality antibody–antigen interfaces occur across diverse epitope geometries.

### PyRosetta-extracted physical features

We use PyRosetta (Chaudhury et al., 2010) to compute a set of features characterizing the physical properties of the antibody–antigen interface, individual chains, and the overall complex. These include measures of interface geometry (e.g., shape complementarity, buried surface area), binding energetics (e.g., Δ*G* and energy decomposition terms), and interaction-specific features such as hydrogen bonding, steric repulsion, and solvation effects.

As shown in Table 4, these Rosetta-derived features exhibit meaningful correlations with DockQ, highlighting the importance of physical interaction quality in determining successful antibody–antigen docking. Metrics associated with interface geometry and binding strength, such as shape complementarity and binding free energy (Δ*G*), show the strongest correlations with DockQ, indicating that well-packed and energetically favorable interfaces are more likely to be predicted accurately. Steric repulsion terms (e.g., fa_rep) are strongly negatively correlated, reflecting the detrimental effect of clashes on interface quality, while van der Waals attraction and hydrogen bonding features show positive correlations, consistent with their role in stabilizing native-like interactions. In contrast, many global or single-chain features exhibit weaker and less consistent correlations, particularly in per-PDB analyses, suggesting that interface-specific properties are more predictive of docking success than overall structural quality. The results of Table 4 is run on the same dataset as Table 2 allowing for direct comparisons of physics based features to confidence metrics. Overall, these results indicate that physically grounded interaction features remain strongly aligned with model performance, even when computed from predicted rather than experimental structures.

### Antigen structural similarity

At the dataset level, we constructed a Foldseek (Van Kempen et al., 2024) database of antigen structures from all 3,800 SAbDab entries and performed an all-by-all structural comparison with a coverage threshold of 0.8. We then clustered antigens across a range of TM-score thresholds. Although the dataset contains 3,800 complexes, this corresponds to only 3,345 unique antigen sequences due to repeated antigens across different antibody complexes. At moderate TM-score thresholds, the number of clusters collapses to fewer than 500, indicating substantial structural redundancy and limited fold diversity among certain antigens. (Figure 13).

**Figure 13:**
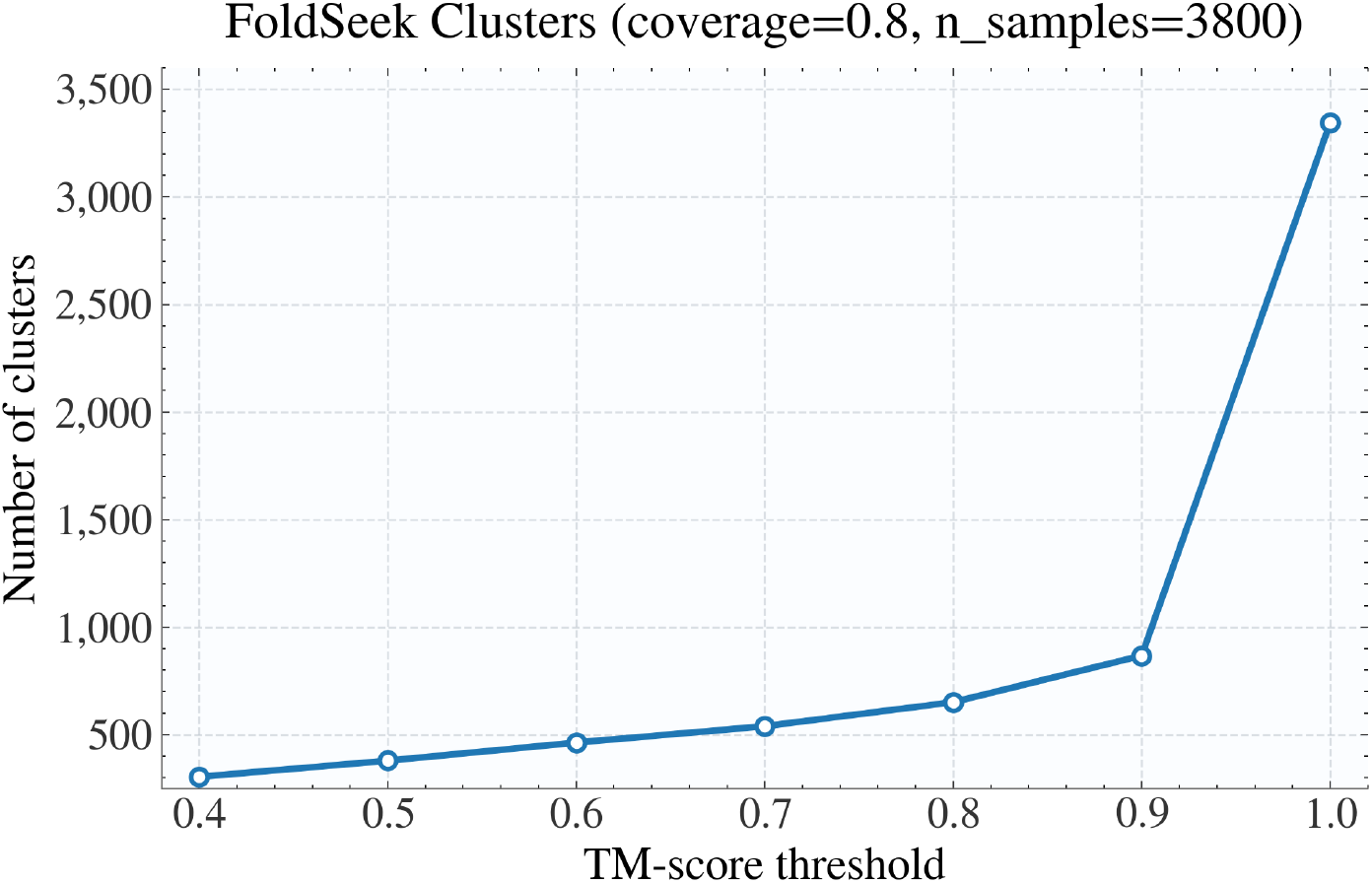
Antigen structural diversity across Foldseek clustering thresholds. Although the dataset contains 3800 complexes, this corresponds to only 3345 unique antigens due to repeated antigens across complexes. Clustering at a TM-score threshold of 0.6 reduces these to 463 structural clusters, indicating substantial redundancy at the fold level.

## Notes

### Competing Interest Statement

D.J.D. owns Intelligent Proteins LLC where he consults biotechnology companies on AI protein engineering. D.J.D. is a co-founder of Metabologic AI, which focuses on developing commercial enzymes with AI. Other authors declare no competing interests.

